## Supplementary #1 for "BOLD Moments: modeling short visual events through a video fMRI dataset and metadata"

### Supplementary #1 Material

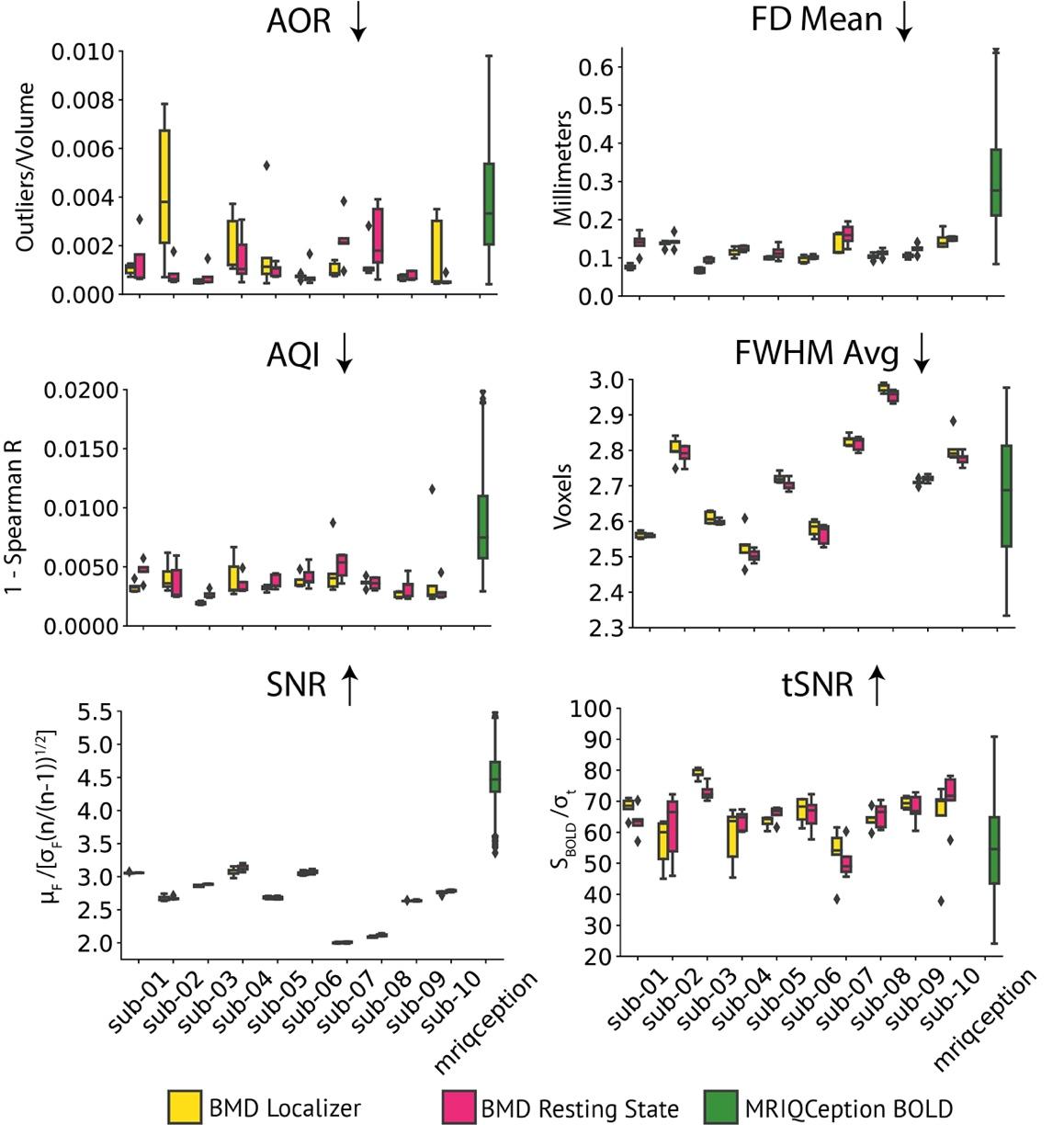

#### **Figure S1: Localizer and resting state functional scan data quality**

Panels compares boxplots of IQM values for each subject for the localizer and resting state functional tasks in the BOLD Moments experiment (per subject: localizer, n=5; resting state, n=5) with anonymous BOLD data pulled from the MRIQCeption API (BOLD, n=687). MRIQCeption does not distinguish between different tasks within BOLD scans. The boxplot extends 1.5 times the high and low quartiles, with outliers defined as a scan with a value outside that range and denoted by diamonds. The up or down arrows after the IQM title correspond to whether higher or lower IQM values denote higher data quality. X-axis labels are shared vertically.

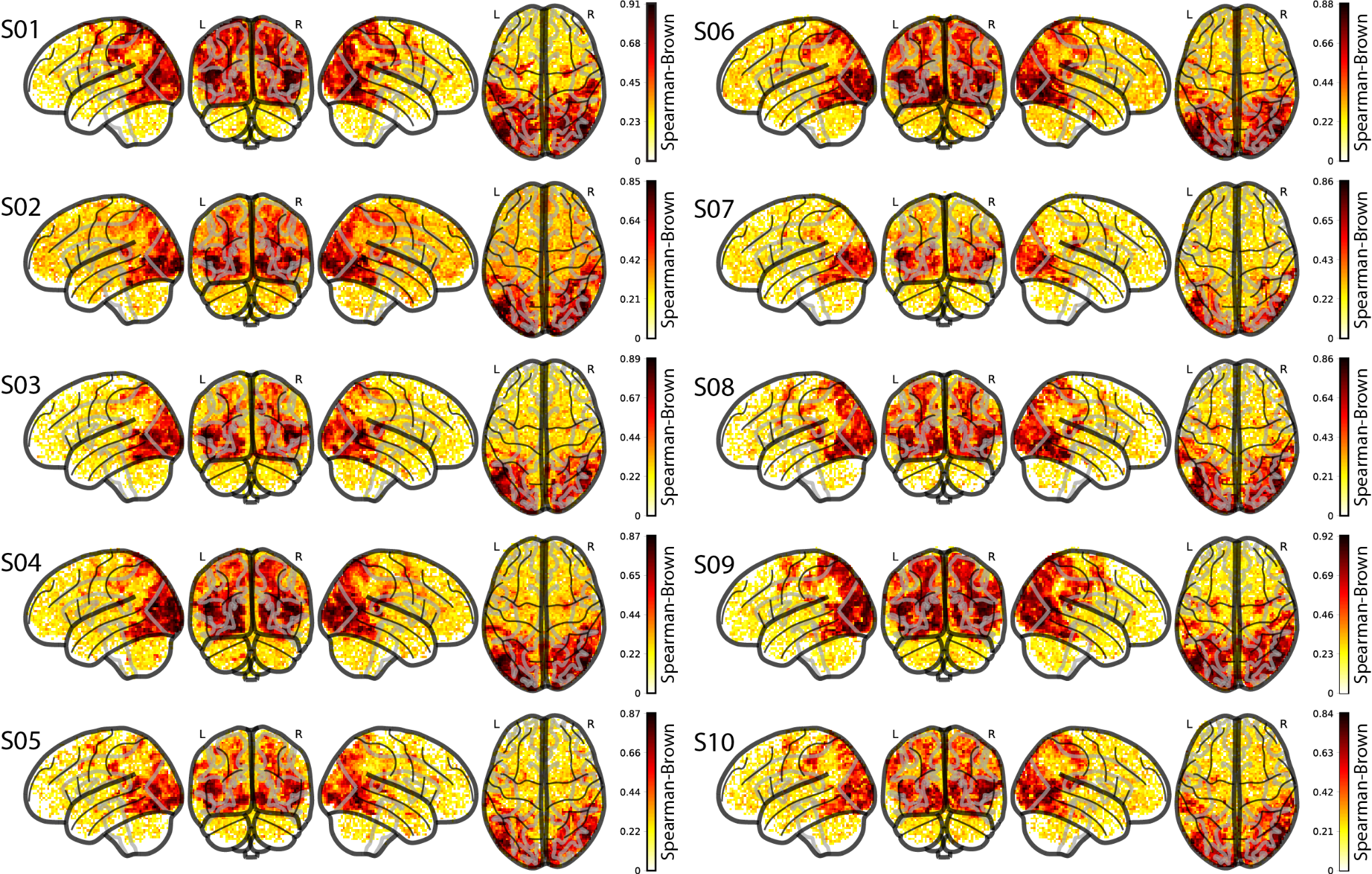

#### **Figure S2: Whole-brain split-half reliability for all subjects**

The glass brains show the split-half reliability (Spearman Brown) at every voxel for each of the ten subjects. A Pearson R correlation value was obtained by correlating random splits of the 10 repetitions from the 102 testing videos. The Spearman Brown split-half reliability was computed using the Pearson R ($\rho$) value obtained above in the formula: Spearman Brown = ($2\rho/(1+\rho)$).

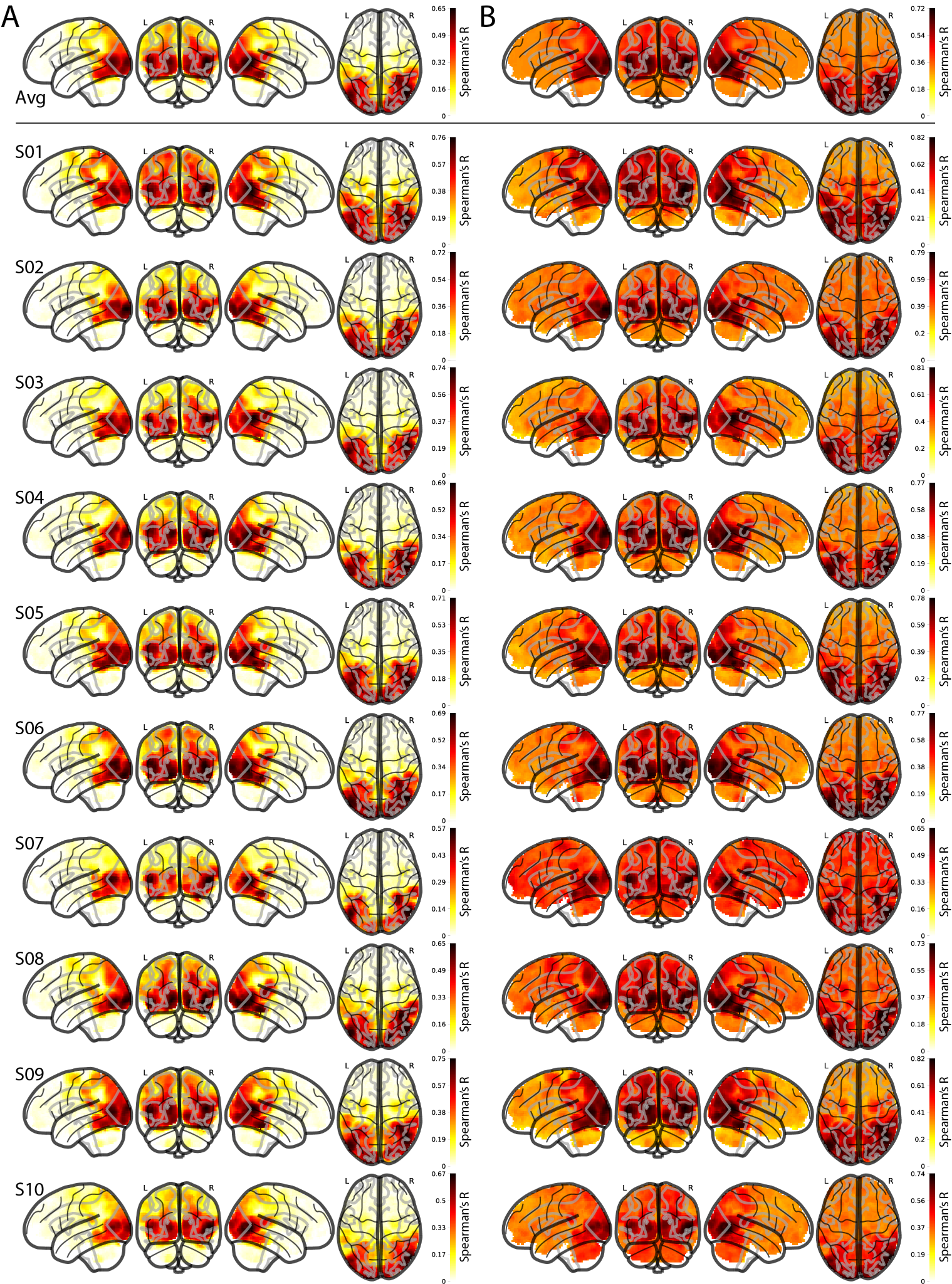

#### **Figure S3: Whole-brain multivariate searchlight-based noise ceilings for all subjects**

**(A) Lower noise ceiling:** The lower noise ceiling is computed using a leave-one-out correlation procedure, where a subject’s RDM at a given voxel *v* is correlated (Spearman’s R) with the remaining nine-subject average RDM at that voxel *v*, repeated over all voxels. **(B) Upper noise ceiling:** The upper noise ceiling is computed by correlating (Spearman’s R) a subject’s RDM at a given voxel *v* with the ten-subject group average RDM at that voxel *v*, repeated over all voxels. We show the whole-brain visualization for the upper and lower noise ceilings averaged over all subjects (top row) and each subject individually (bottom). The brain responses used to compute the RDMs are from the beta values of the testing set.

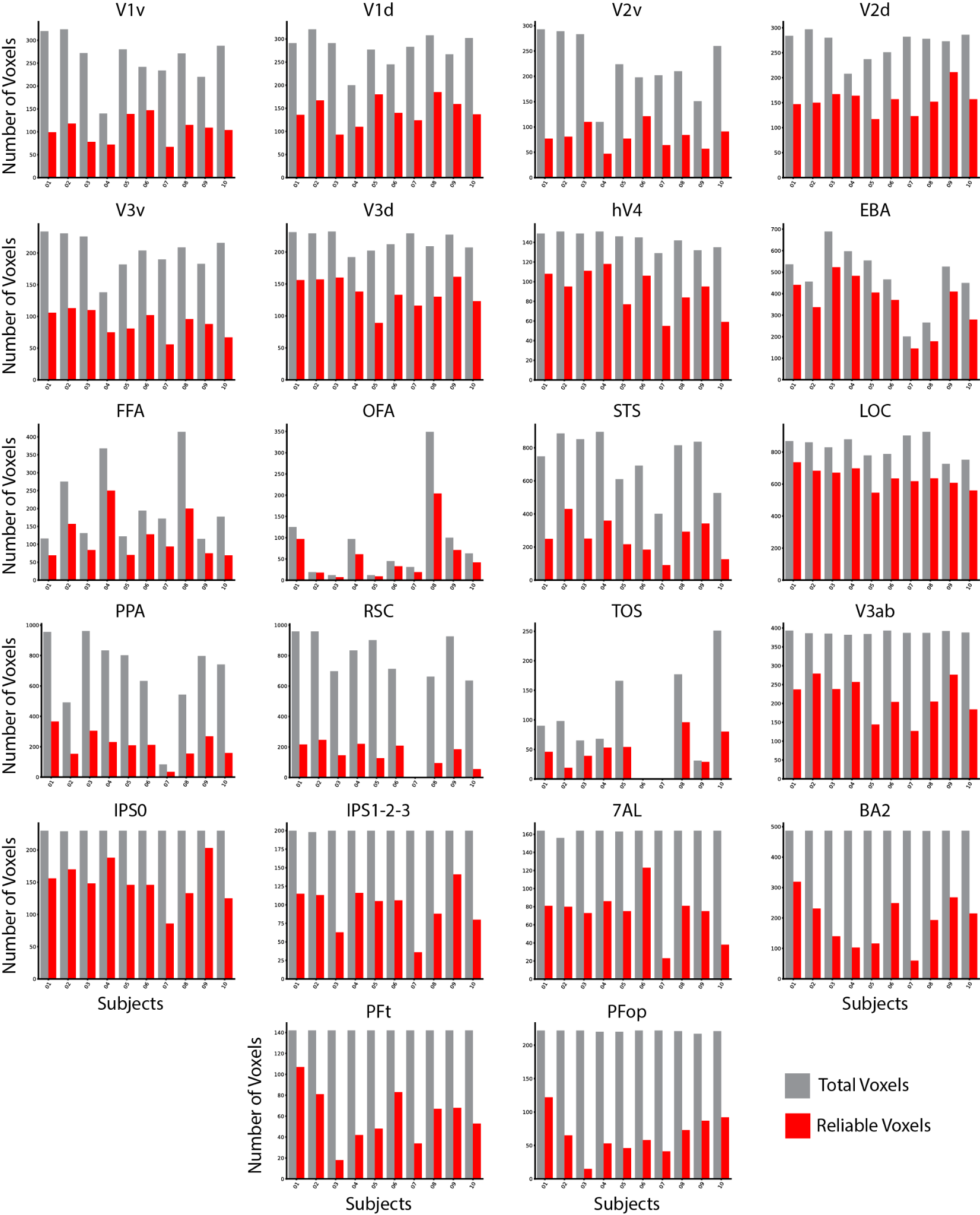

#### **Figure S4: ROI reliability and size separated for each subject**

Barplots of the total number of voxels in the ROI mask (gray bar) and the total number of reliable voxels (p < 0.05, Spearman-Brown) in the ROI mask (red bar) for each subject across the twenty-two ROIs. Subject 6 did not show any activation from the functional localizer task for ROI TOS, and subject 7 did not show any activation from the functional localizer task for ROIs RSC and TOS. Y-axis and X-axis labels are shared horizontally and vertically, respectively.

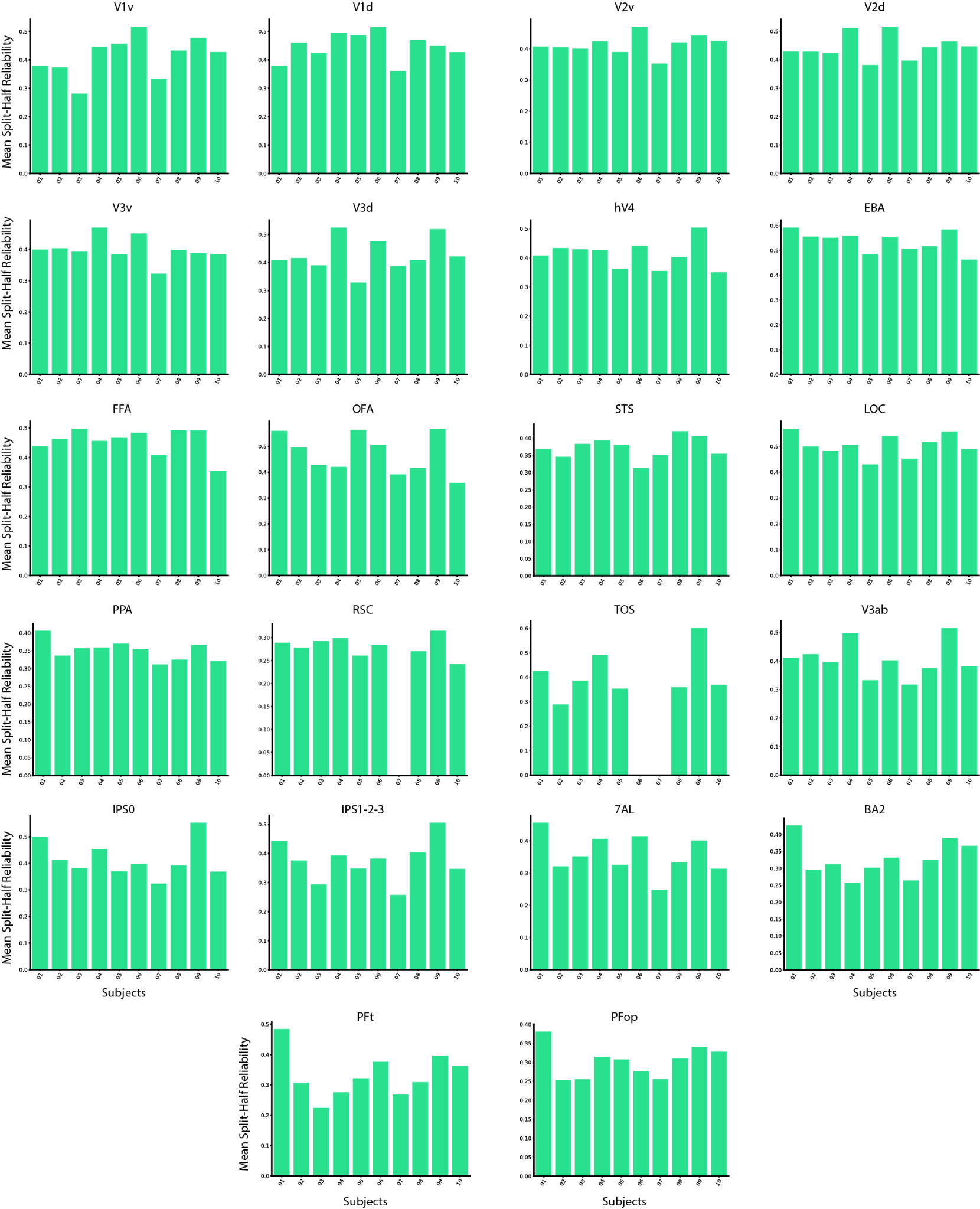

#### **Figure S5: Average reliability in each ROI for each subject**

Barplots of the average split-half reliability in each ROI of all reliable voxels, separated for each subject. Y-axis and X-axis labels are shared horizontally and vertically, respectively.

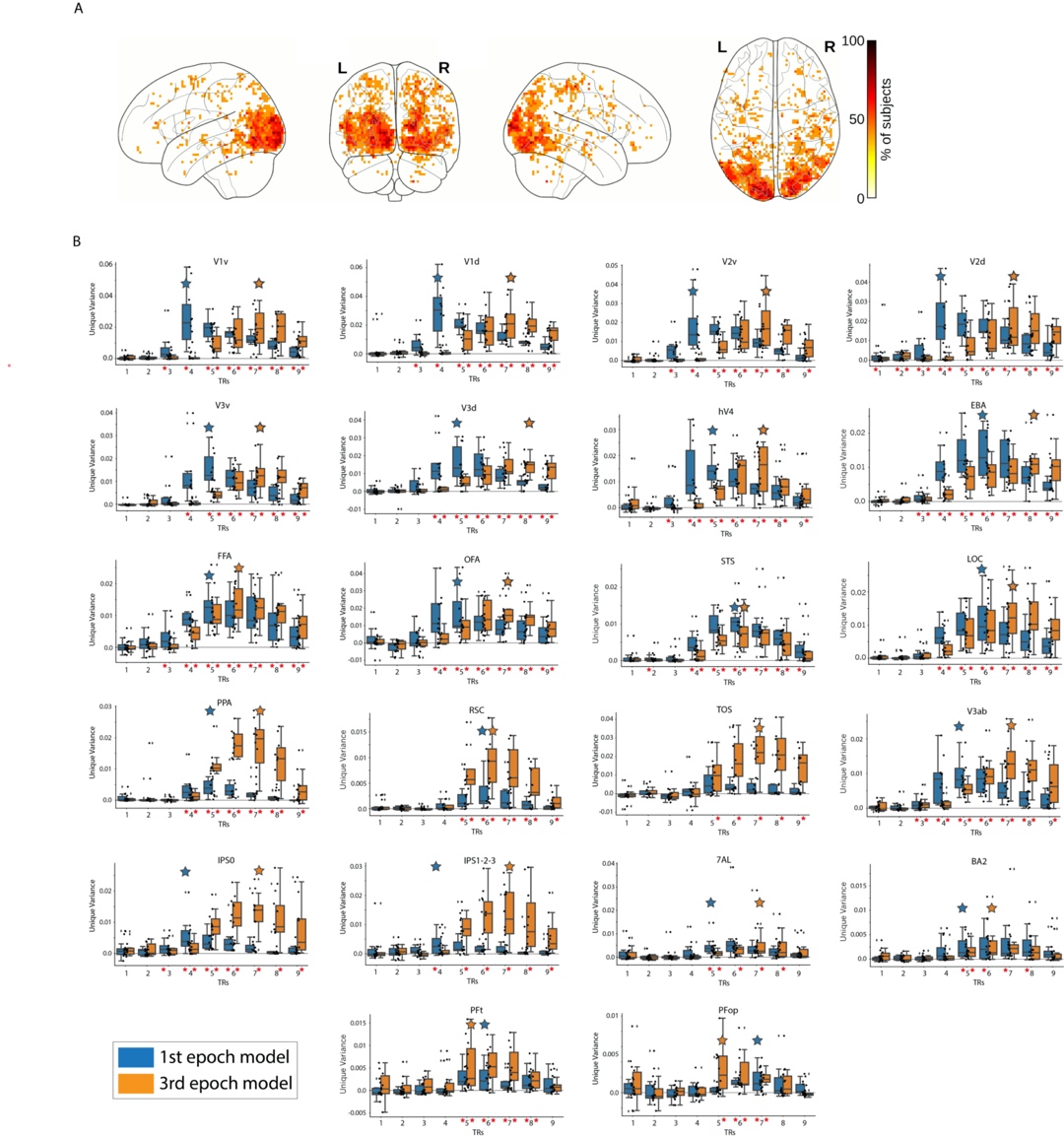

#### **Figure S6: Encoding the temporal dynamics of the BOLD signal.**

**(A)** **Whole-brain analysis:** Each voxel shows the percentage of subjects with a TR peak difference of 2 TRs at that specific voxel. Only significant voxels are plotted (p < 0.05, binomial test, FDR corrected). The effect of interest is showing predominantly in the visual cortex. **(B) ROI analysis:** Unique variance explained by the first and third video epoch (second) synthetic fMRI data, at each TR. Red asterisks along the x-axis indicate unique variance scores significantly greater than 0 (p<0.05, one-sample one-side t-test, FDR corrected across 9 TRs x 2 video epochs = 18 comparisons). Large blue/orange stars indicate the TR with the highest subject averaged unique variance for the first/third video epochs, respectively. The box plot encompasses the first and third data quartiles and the median (horizontal line). The whiskers extend to the minimum and maximum values within 1.5 times the interquartile range, and values falling outside that range are considered outliers (denoted by a diamond). The overlaid points show the value at each observation (n=10 for all ROIs except TOS (n=8) RSC (n=9)). Y-axis and X-axis labels are shared horizontally and vertically, respectively.

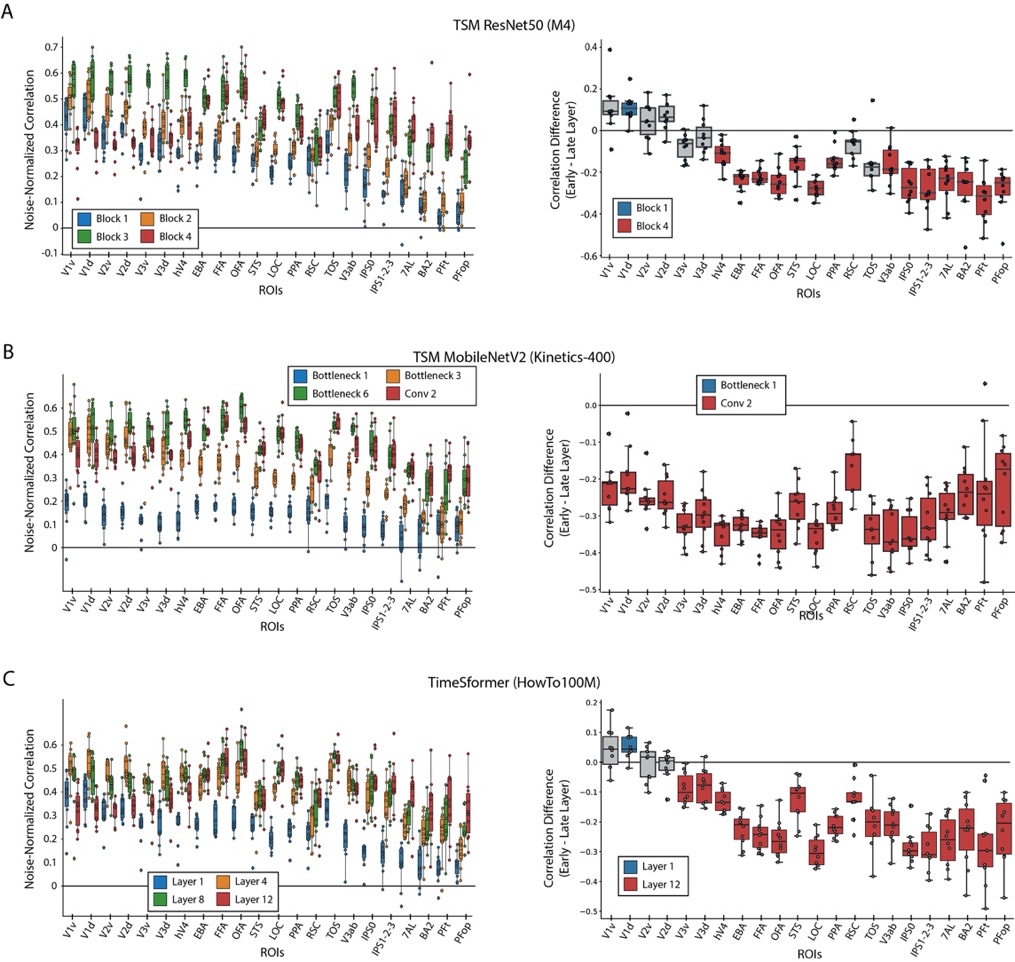

#### **Figure S7: Encoding model performance on BMD**

A) TSM ResNet50 trained on M4: Features were extracted after block’s 1 (blue), 2 (orange), 3 (green), and 4 (red) in the ResNet 50 architecture. B) TSM MobileNetV2 trained on Kinetics-400: Features were extracted after the first bottleneck layer (blue), third bottleneck layer (orange), sixth bottleneck layer (green), and last 2D convolutional layer before the average pool (red) in the MobileNetV2 architecture. C) TimeSformer S+T trained on HowTo100M: Of the model’s twelve layers, features were extracted after the first (blue), fourth (orange), eighth (green), and twelfth (red) layers. The box plot on the left side in each panel shows the noise-normalized predictivity of four of each architecture’s features at each of the 22 ROIs. The features were extracted at early (blue), intermediate (orange and green), and late (red) processing stages in each architecture to capture increasingly high-level degrees of transformations. The box plot on the right side in each panel shows the brain prediction difference between each architecture’s latest and earliest layers for each subject and ROI. For the box plots on the right, a blue or red colored box plot denotes a significant difference in correlations from 0 (p<0.05, two-sided one-sample t-test, Bonferroni corrected for n=22 comparisons), and gray denotes no significance. The box plots encompass the first and third data quartiles and the median (horizontal line). The whiskers extend to the minimum and maximum values within 1.5 times the interquartile range, and values falling outside that range are considered outliers (denoted by a diamond). The overlaid points show the value at each observation (n=10 for all ROIs except TOS (n=8) RSC (n=9)).

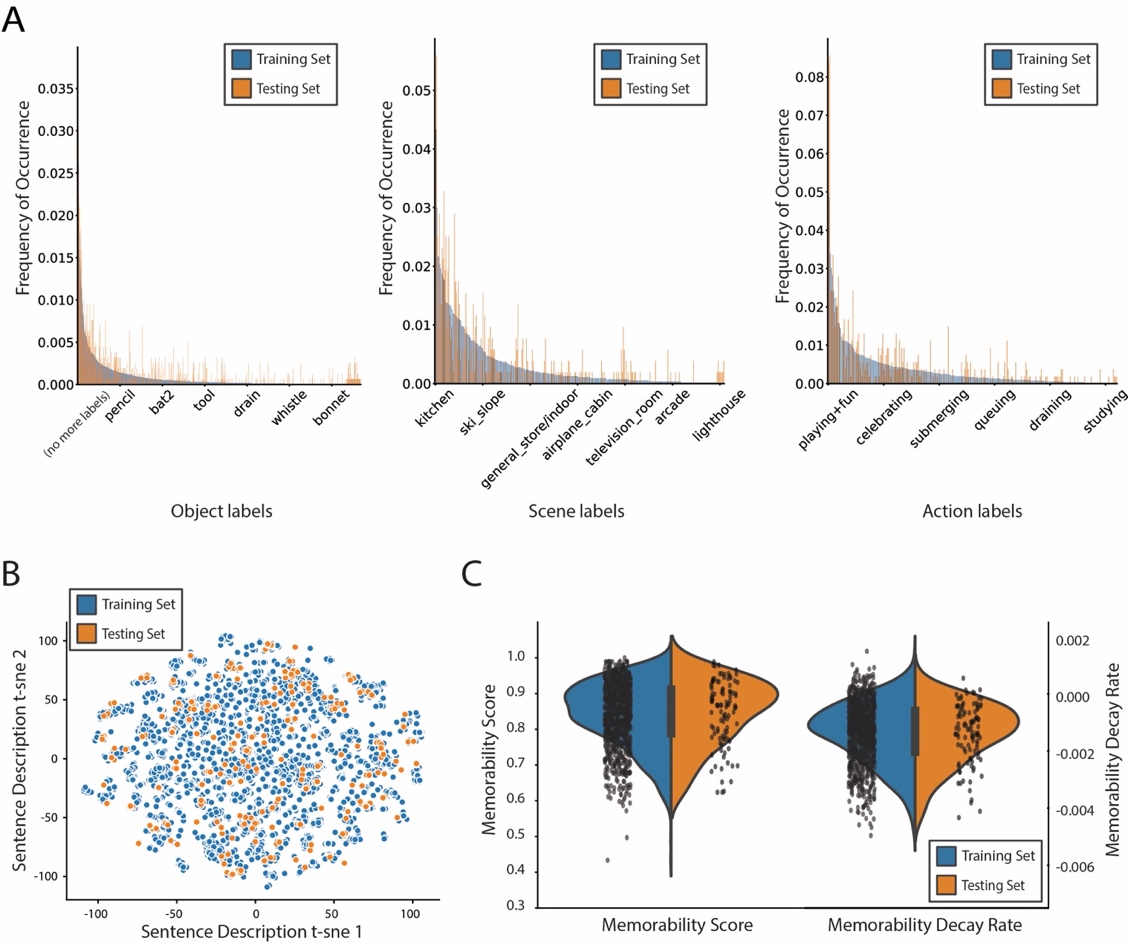

#### **Figure S8: Distributions of stimuli metadata between training and testing sets**

The training and testing sets consist of 1,000 and 102 different videos, respectively. An author manually inspected the pairs of testing set videos to ensure no high-level semantic overlap, in terms of objects and actions. **A) Object, scene, and action label frequency of occurrence:** The barplot depicts the frequency of occurrence, (between 0 and 1) of, from left to right, the single-word object, scene, and action labels of the 1,102 video stimuli used in the BOLD Moments Dataset. The frequency bars for each label are separated by training (blue) and testing (orange) splits to show their similar frequency of distributions. **B) Text description and spoken transcription t-sne distances.** The scatterplot shows the t-sne components (n=2 components, perplexity=10, number of iterations=1000) of each text description or spoken transcription embedding. The 6 sentence descriptions per video (5 text descriptions and 1 spoken transcription) serve as a useful proxy for the video’s content. The t-sne plot shows the training and testing set stimuli cover similar spaces of video content. **C) Memorability distribution.** The distribution of the memorability scores and memorability decay rates (1 per video) between the training and testing splits are highly similar and approximately normal. Note that the positive memorability decay rates, while theoretically implausible, reflect the true experimental results detailed in the Memento10k dataset. Users may want to set positive values to 0 depending on the analysis.

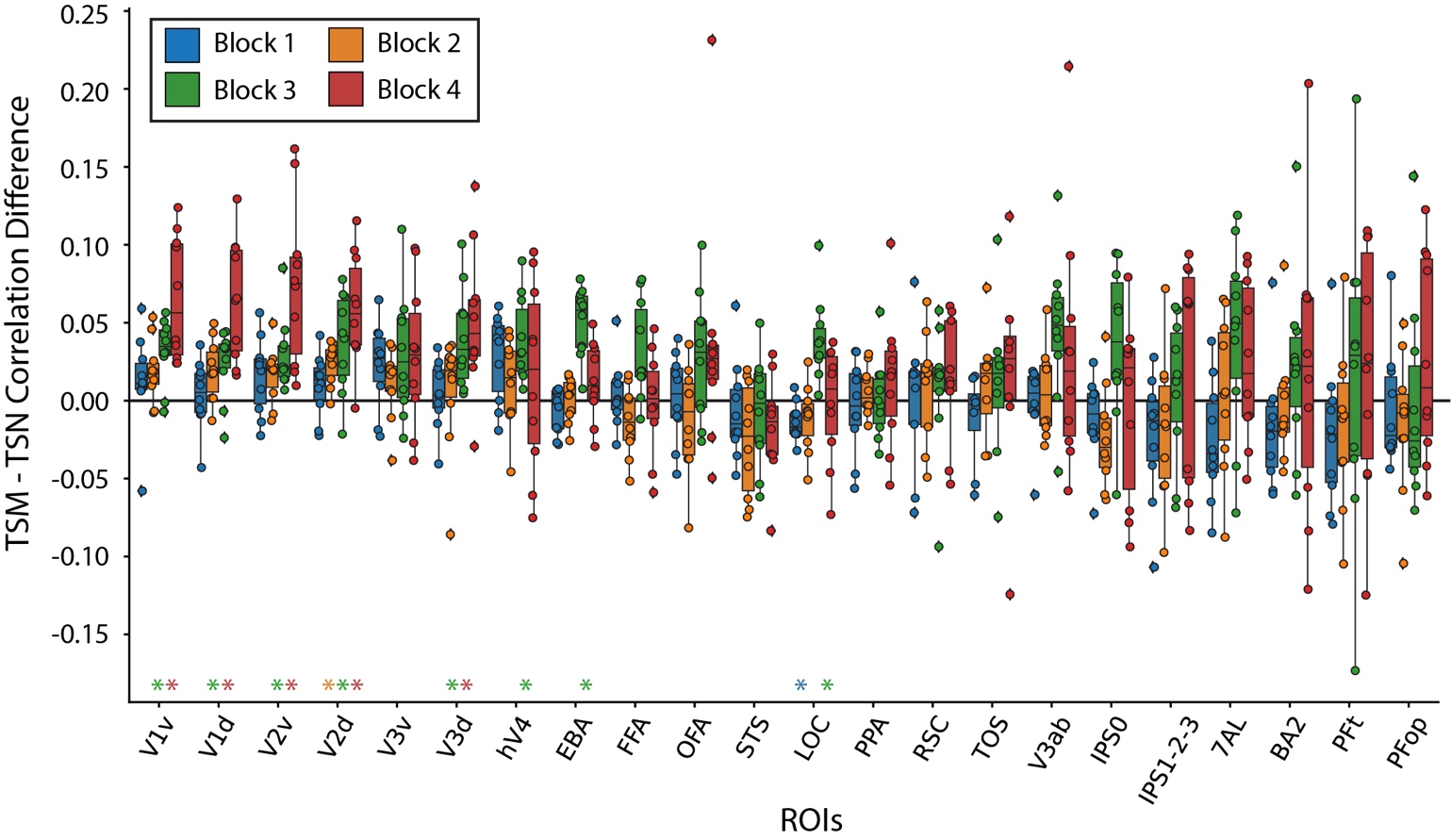

**Figure S9: The effect of Temporal Shift Module (TSM) on brain prediction performance**

**(A) TSM vs TSN prediction performance:** The difference in subject brain prediction performance of a TSM ResNet50 and Temporal Segment Network (TSN) ResNet50 each trained on a 10,000-video subset of the M4 dataset (Multi-moments Minus Memento) was computed at each of the four Blocks for each ROI. TSM results in increased brain prediction performance most prominently in early visual ROIs. In both panels, colored asterisks along the x-axis plot indicates significant difference between the unshuffled and shuffled prediction accuracy at that DNN block (one sample two-sided t-test against a population mean of 0, FDR correction across 22 ROIs x 4 blocks = 88 comparisons, p < 0.05). The box plot encompasses the first and third data quartiles and the median (horizontal line). The whiskers extend to the minimum and maximum values within 1.5 times the interquartile range, and values falling outside that range are considered outliers (denoted by a diamond). The overlaid points show the value at each observation (n=10 for all ROIs except TOS (n=8) RSC (n=9)).

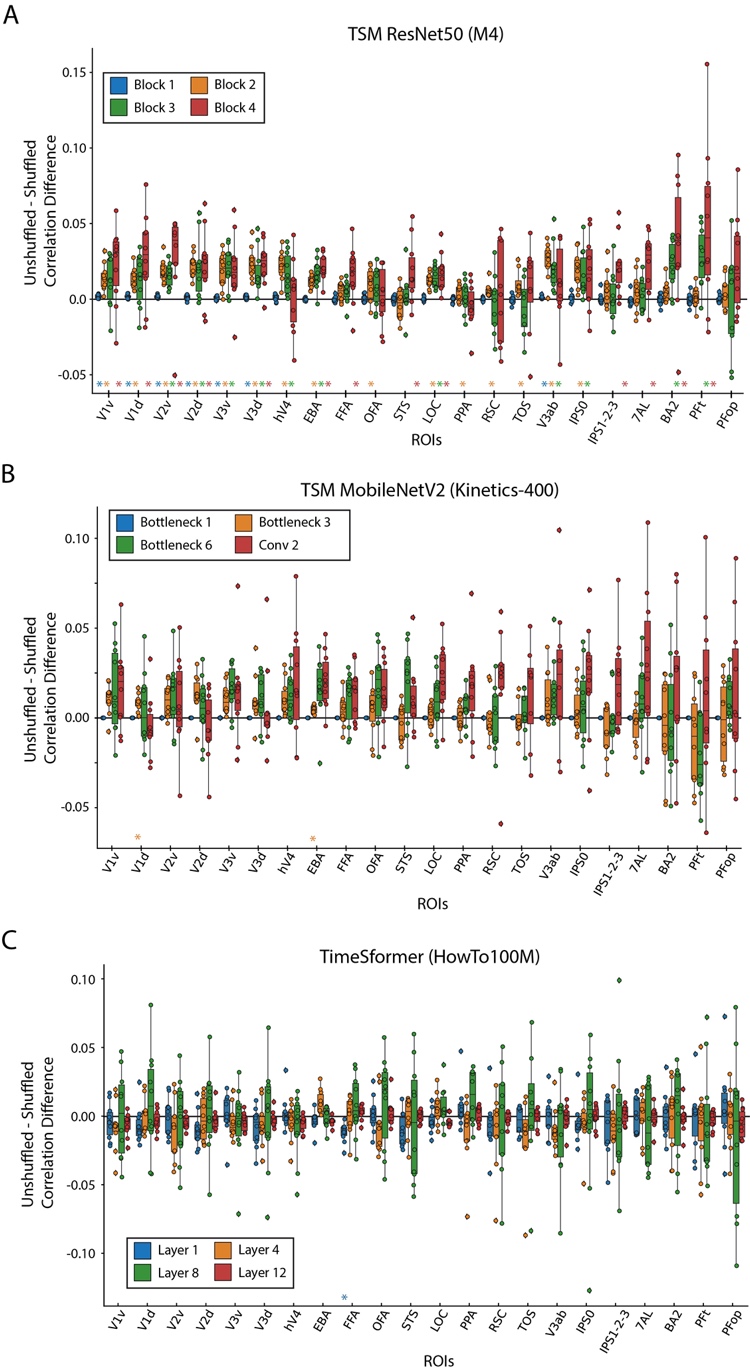

**Figure S10: The effect of frame shuffling on brain prediction performance across different architectures**

We compute the difference in the correlation between the shuffled frame prediction accuracy and unshuffled frame prediction accuracy at all 22 ROIs and four layers of a **(A)** TSM ResNet50, **(B)** TSM MobileNetV2, and **(C)** TimeSformer model. Features were extracted at increasing levels of depth in each model (blue, orange, green, red) that reflect higher levels of model processing stages. Only the TSM ResNet50 architecture trained on the M4 dataset (Multi-moments Minus Memento10k) showed evidence of robust differences across cortex between shuffled and unshuffled input. Colored asterisks along the x-axis plot indicates significant difference between the unshuffled and shuffled prediction accuracy at that DNN block (one sample two-sided t-test against a population mean of 0, FDR correction across 22 ROIs x 4 blocks = 88 comparisons, p < 0.05). The box plot encompasses the first and third data quartiles and the median (horizontal line). The whiskers extend to the minimum and maximum values within 1.5 times the interquartile range, and values falling outside that range are considered outliers (denoted by a diamond). The overlaid points show the value at each observation (n=10 for all ROIs except TOS (n=8) RSC (n=9)).

| **Original Citation** | **Dataset Name** | **Number of Subjects** | **Number of Unique stimuli**  **(Shared across all subjects)** | **Stimulus Repetitions Within Subject**  **(Stimuli x repetitions)** | **Provided Stimulus Metadata?** | **Stimulus Superset** | **MRI Scanner Strength of Main Task** | **Auxiliary Measurements (in addition to structural)?** | **Other Neuroimaging Modalities** | **Experiment Superset** |
| --- | --- | --- | --- | --- | --- | --- | --- | --- | --- | --- |
| Ours | BOLD Moments | 10 | 1,102 3 second videos (1102) | 1,000 videos x 3  102 videos x 10 | Yes | Moments in Time  Multi-Moments in Time  Memento10k | 3T | Yes | EEG (in progress) | None |
| (Chang et al., 2019) | BOLD5000 | 4 | 4,916 images (113)^a^ | 4,916 images x 1 | Yes | SUN  COCO  ImageNet | 3T | Yes | None | None |
| (Hanke et al., 2016) | Forrest Gump | 15 | 1 2-hour movie (1) | 1 movie x 1 | Yes | None | 3T | Yes | fMRI (audio-only)^c^  fMRI (music listening) | StudyForrest |
| (Allen et al., 2022) | NSD | 8 | 70,566 images (515) | 10,000^d^ images x 3 | Yes | COCO | 7T | Yes | None | None |
| (Seeliger et al., 2019) | Doctor Who | 1 | 30 45-minute TV episodes (n/a)  7 1-3 minute video clips (n/a) | 30 episodes x 1  7 video clips x 22 | No | None | 3T | Yes | None^e^ | None |
| (Aliko et al., 2020) | Naturalistic Neuroimaging Database (NNDb) | 86 | 10 movies (10)^f^ | 1 movie x 1 | No | None | 1.5T | Yes | None^g^ | None |
| (Hebart et al., 2023) | THINGS-data | 3 | 8,740 images (8,740) | 8640 x 1  100 x 12 | Yes | THINGS | 3T | Yes | MEG  EEG | THINGS initiative |
| (Nishimoto et al., 2011) | Vim-2 | 3 | 2 continuous video streams (2) | 1 stream (7200 seconds) x 1  1 stream (540 seconds) x 10 | No | None | 4T | No | None | None |
| (Wen et al., 2018) | n/a | 3 | 2 continuous video streams (2) | 1 374-clip stream (2.4 hours) x 2  1 598-clip stream (40 minute) x 10 | No | None | 3T | No | None | None |
| (Zhou et al., 2023) | Human Action Dataset (HAD) | 30 | 21,600 2 second video clips (0) | 720 videos x 1 | Yes | Human Action Clips and Segments (HACS) Clips | 3T | No | MEG (in progress) | None |
| (Boyle et al., 2020) | Friends s01 - s06 | 6 | 146 22-minute TV episodes (146)^h^ | 1 episode x 1 | No | None | 3T | Yes | None | Courtois NeuroMod |
| (Boyle et al., 2020) | movie10 | 6 | 4 ~1-3 hour movies (4) | 2 movies x 1  2 movies x 2 | No | None | 3T | Yes | None | Courtois NeuroMod |
| (Horikawa & Kamitani, 2017) | Generic Object Decoding (GOD) | 5 | 1,250 images (1,250) | 1200 images x 1  50 images x 35 | Yes | ImageNet | 3T | Yes | fMRI (mental imagery) | None |

^a^ Chang et al., 2019: 4 subjects saw 112 images, 3 subjects saw 1 image

^b^ Chang et al., 2019: Images were sampled from SUN (1000), COCO (2000), and ImageNet (1916)

^c^ Hanke et al., 2016: Audio-visual fMRI was originally acquired

^d^ Allen et al., 2022: Subjects each saw between 9-10,000 images

^e^ Seeliger et al., 2019: Audio-visual fMRI was originally acquired

^f^ Aliko et al., 2020: 8 movies were seen by 6 subjects, 1 movie was seen by 18 subjects, and 1 movie was seen by 20 subjects

^g^ Aliko et al., 2020: Audio-visual fMRI was originally acquired

^h^ Boyle et al., 2020: Subject 04 completed seasons 1-4 and part of season 5

#### Table S1: Comparison of large visual fMRI datasets

We compare large, naturalistic, task-based, visual fMRI datasets across features of interest to computational neuroscientists for modeling purposes. The features highlight a dataset’s stimuli (type, number, overlap with existing stimuli sets, and annotations), fMRI acquisition (scanner strength, auxiliary measurements, complementary neuroimaging modalities, and subset in greater neuroimaging efforts), and experimental design (number of subjects and stimulus repetitions). This table summarizes a niche each dataset fills and should not be mistaken for a comparison of dataset quality. Values may differ slightly from the original publication in order to facilitate comparisons across datasets and necessarily summarize information. Note that while some datasets are not officially part of a larger experiment superset, many have been used in independent studies and thus may have additional stimuli metadata and neuroimaging data. Such cases are not noted in this table to maintain clarity. Please see the original publication for the most accurate information.

The added value of brain responses to a short video dataset versus a static image dataset

We emphasize that a short video (e.g., 3 second duration, as in BMD) fMRI dataset is not better or worse than a static image fMRI dataset; rather, they are different in terms of stimulus features and corresponding brain responses that may make one better suited to answer specific research questions. Most obvious, short videos contain a temporal dimension that static images do not, allowing the video to communicate crucial contextual information about how spatial components in our environment move (or not) and spatially relate to each other over time. The benefit of this temporal dimension is clear in our everyday lives – we can interpret transitions between states (a door is being opened, not closed), direction (a steering wheel is being turned to left, not right or still), reactions (the child laughed when shown the picture), motion (the baby is crawling slowly, not fast), and more.

The contextual value of a video’s temporal dimension is reflected in BMD’s own action and sentence text description metadata. Concerning action labels, images can only be labelled with a limited subset of actions or else be highly constrained in order to capture a specific action. For example, the action of a baseball player “hitting” the ball can only be captured with an image if the photo were taken at very specific instant in time, otherwise the action may be ”standing” or “swinging”. Even a short video like in BMD easily captures these actions without heavily constraining the space of possible videos that correspond to “hitting”. Concerning text descriptions, short videos can capture temporal sequences of events that an image cannot. Examples of such captions sampled from some of BMD’s first videos include (emphasis our own):

- Video 0001: "A mallard is in the water alone *swimming around* and *putting its beak in.*"
- Video 002: "A man *is showing another man* how to *move feet back and forth*."
- Video 005: "A woman guides a little boy's arms *up and down as other kids stretch* around him."
- Video 006: "a chess tournament is going on this is focused on two players one *is moving their queen and taking something* to put the king in checkmate"

Static frames of these videos cannot capture the temporal facts that the mallard is “putting its beak in”, the man “is showing another man how”, “a woman guides…as other kids stretch”, and a chess player “is moving their queen and taking something to put the king in checkmate.” This temporal information adds valuable context that often makes one’s understanding of the 3s video vastly different compared to any one of its single static frames.

But do these differences in short videos and static images translate to differences in fMRI brain responses? Yes, previous work has found that videos evoke a greater extent (Bartels & Zeki, 2004; Konen & Kastner, 2008; Press et al., 2001; Schultz & Pilz, 2009; Yildirim et al., 2019) and pattern (Buccino et al., 2004; Kret et al., 2011; Lingnau & Downing, 2015; Wurm & Caramazza, 2022) of cortex responding to videos than images throughout occipitotemporal, dorsal visual, and parietal cortex. In this manuscript we describe our highly reliable activations throughout cortex (Figure 3) with notably high reliability in parietal cortex, a region of the brain that weakly responds to static images. These highly reliable brain responses are not just a result of increased participant engagement or stimulus saliency; we even show that BMD brain responses capture temporal information from the videos (Figure 5, Figure 6, Supplementary 1 Figure 9, Supplementary 2 Figure 2) despite the BOLD response’s temporal sluggishness and fMRI’s low sampling rate.

In the neighboring field of computer vision, researchers have long recognized that videos and images demand different modeling approaches (Ahn et al., 2023; Bertasius et al., 2021; Lin et al., 2019; Tong et al., 2022; Wang et al., 2016) and training datasets (Goyal et al., 2017; Kay et al., 2017; Miech et al., 2019; Monfort et al., 2020; Soomro et al., 2012) for strong task performance. Videos continue to be at the forefront of ground breaking computer vision research due to their creative, cross-domain, and practical applications in text-to-video generation (Ho et al., 2022; Singer et al., 2022; Wu et al., 2023), video understanding with large language models (Ju et al., 2022; Maaz et al., 2023; Zhang et al., 2023), and efficient action recognition and pose estimation (Liu et al., 2023; Qing et al., 2024; Zheng et al., 2023).

Taken together, short video fMRI datasets offer unique opportunities to advance the field of computational neuroscience where static image fMRI datasets cannot. They can advance methodologies around estimating BOLD signals in response to rapid stimulus presentations (Misaki et al., 2013; Prince et al., 2022; Wittkuhn & Schuck, 2021), elucidate cognitive functions concerning temporal integration (Fairhall et al., 2014; Hasson et al., 2008; Orlov & Zohary, 2018), test temporally specific cognitive objective functions (Doerig et al., 2022; Kanwisher et al., 2023), and detail how multiple visual pathways interact to achieve an understanding of an event (Lingnau & Downing, 2015; Mineault et al., 2021; Pitcher & Ungerleider, 2021; Wurm & Caramazza, 2022). As neuroscience and computer science research become increasingly intertwined (Allen et al., 2022; Chen et al., 2023; Cichy et al., 2019, 2021), BMD is well-suited to integrate with state-of-the-art video modeling work from the computer vision community. Importantly, a short video dataset like BMD can make these scientific advancements while staying connected to the vast body of still image work by sharing event-related paradigms, multivariate and univariate methodologies, representational similarity analyses, and/or encoding and decoding techniques. Short video datasets offer more ecological validity than static images while retaining experimental control and offer tremendous potential to advance our understanding of the human visual system.

#### Structural and functional scan quality assessment

We use MRIQC (Esteban et al., 2017) to measure the quality of our study’s original or minimally preprocessed structural and functional MRI scans. MRIQC is an open-source software that outputs a large and diverse set of image quality metrics (IQMs) to comprehensively quantify the quality of (f)MRI data in a standardized and reproducible manner. IQMs are calculated at the level of a single run, and group reports are generated for all T1w, T2w, and BOLD runs in the study. We present a representative subset of 6 IQMs to summarize the quality of our structural scans and another subset of 6 IQMs to summarize the quality of our functional scans (see MRIQC documentation for details on all 112 IQMs:  [https://mriqc.readthedocs.io/en/latest/measures.html](https://www.google.com/url?q=https://mriqc.readthedocs.io/en/latest/measures.html&sa=D&source=docs&ust=1700961676876197&usg=AOvVaw2tB9D6uITvdnuysrYQtTlw)). Note that no set of metrics can fully describe data quality in itself. Thus, when choosing IQMs to represent the structural and functional scan quality, we primarily considered the following three criteria:

First, the representative IQMs for the structural scans and for the functional scans should capture metrics especially relevant to the properties of structural and functional scans.

Second, IQMs that are useful for describing the quality of both structural and functional scans are preferred in order to create more cohesive and shared IQM subsets between the structural and functional scans.

Third, IQMs commonly reported in previous literature are preferred in order to increase comparisons across studies and be more familiar to readers.

We additionally use MRIQCeption to contextualize our study’s group reported results within a large collection of anonymized group reports from studies of comparable scanner parameters (1 < Tesla < 3, 1 <= TR < 3).

For structural (T1w and T2w) scans, we present the results from the following IQMs:

**SNR Total - Signal to Noise Ratio:** SNR Total for structural scans is computed by averaging the SNR across the cerebrospinal fluid (snr_csf), gray matter (snr_gm), and white matter (snr_wm). SNR is calculated by the following formula:

$$\frac{\mu_{F}}{\sigma_{F}\sqrt{n(n-1)}}$$

Where $\mu_{F}$ is the mean intensity of the foreground, $\sigma_{F}$ is the standard deviation of the foreground intensity, and $n$ is the number of voxels in the foreground mask. Higher values correspond to higher quality.

**CNR - Contrast to Noise Ratio:** CNR, an extension of SNR, computes the absolute value difference of the gray and white matter image values (|S_W_ - S_G_|) and divides them by the standard deviation of the values in the surrounding air (σ_air_). Higher values correspond to higher quality.

**CJV - Coefficient of Joint Variation:** CJV is the ratio of the coefficient of variation in the gray matter to the coefficient of variation in the white matter. Lower values correspond to higher quality.

**EFC - Entropy Focus Criterion:** EFC is the shannon entropy of voxel intensities normalized by the maximum shannon entropy value. It measures ghosting and blurring due to head motion. Lower values correspond to higher quality.

**FWHM Avg - Average Full-Width Half Maximum Smoothness:** FWHM Avg is the average spatial distribution of voxel intensities in an image using a gaussian width estimator. Lower values correspond to higher quality.

**FBER - Foreground-Background Energy Ratio:** FBER is the ratio of the mean energy inside the head to the mean energy outside the head. Higher values correspond to higher quality.

For functional scans, we present the results from the following IQMs:

**SNR - Signal to Noise Ratio:** SNR for functional scans is calculated by the following formula:

$$\frac{\mu_{F}}{\sigma_{F}\sqrt{n(n-1)}}$$

Where $\mu_{F}$ is the mean intensity of the foreground, $\sigma_{F}$ is the standard deviation of the foreground intensity, and $n$ is the number of voxels in the foreground mask. Higher values correspond to higher quality.

**tSNR - Temporal Signal to Noise Ratio:** tSNR divides the mean BOLD signal across time by the temporal standard deviation map. Higher values correspond to higher quality.

**FD Mean - Mean Framewise Displacement:** FD Mean computes the average displacement of all six motion parameters. Lower values correspond to higher quality.

**FWHM Avg - Average Full-Width Half Maximum Smoothness:** FWHM Avg is the average spatial distribution of voxel intensities in an image using a gaussian width estimator. Lower values correspond to higher quality.

**AOR - AFNI Outlier Ratio:** AOR is the average fraction of outliers found in each fMRI volume as computed by AFNI’s “3dToutcount” function. Lower values correspond to higher quality.

**AQI - AFNI Quality Index:** AQI computes the average distance between each volume and the median volume of a series, given by AFNI’s “3dTqual” function. Lower values correspond to higher quality.

#### The Algonauts Project 2021 challenge approaches of the top three winners

*The Algonauts Project 2021: How the Human Brain Makes Sense of a World in Motion* is an open challenge that took place during the spring and summer of 2021 and culminated in an interactive workshop and speaking event at the Computational Cognitive Neuroscience (CCN) conference (Cichy et al., 2021; Naselaris et al., 2018). For the challenge, participants submit the predictions of their computational model on held-out brain data (see <http://algonauts.csail.mit.edu/challenge.html> for the final challenge leaderboard and details). We summarize the top three challenge entries, highlighting their different modeling approaches and insights at the intersection of natural and artificial intelligence research.

The first-place team “huze” approached this challenge using an ensemble of 6 different models that together integrate meaningful features of video understanding: spatiotemporal, motion, edge, and audio features (Yang et al., 2021). They then weighted the outputs of each model representation and found that the predictivity for each ROI was highest when combining features from all models. They additionally optimized the receptive field size for each of the four I3D RGB model layers and ROI (Monfort et al., 2020). They showed that early ROIs benefited most from smaller receptive fields on low-level layers (layers 1 and 2) and later ROIs benefited most from larger receptive fields on high-level layers (layers 3 and 4), replicating neuroscience results (Dumoulin & Wandell, 2008).

The second-place team “bionn” was interested in evaluating a range of DNNs from the more classical supervised CNNs (AlexNet, VGG19, ResNet50, and ResNet152) to the more modern contrastive learning and visual transformer networks (simclr, pclv2, and visual transformer network ViT) (Janik & Olesik, 2021). They found the ResNet models, specifically ResNet152, outperformed the visual transformer and contrastive learning networks. Similar to “huze”, “bionn” also took advantage of pooling the model features to simulate small receptive fields for early regions and large receptive fields for later regions.

The third-place team “shinji” (Nishimoto, 2021) experimented with state-of-the-art spatiotemporal vision features from TimeSformer (Bertasius et al., 2021) and classical, neurophysiology-based motion energy features (Nishimoto et al., 2011; Watson & Ahumada, 1985). Looking exclusively at the TimeSformer model, they first saw that earlier layers (layers 4-6 out of 12) best predicted early visual regions (V1-V4) while later layers (layers 9-11 out of 12) best predicted later visual regions (EBA, LOC, STS, FFA, and PPA). In early visual regions (V1-V3), the motion-energy model outperformed the TimeSformer model, and in the later visual regions (V4, EBA, LOC, STS, FFA, and PPA), the TimeSformer model was better. However, the combination of both the TimeSformer and motion-energy features was best for all ROIs except for FFA, STS, and PPA.

For more details about the approaches of the top three challenge winners, see the PDFs of their full reports, available online or with the BMD dataset.
