## Supplementary #2 for "BOLD Moments: modeling short visual events through a video fMRI dataset and metadata"

Supplementary #2 Material

1 Overview

This supplementary file details an additional preprocessed version (version B) of the BOLD Moments Dataset (BMD) released alongside the version presented in the manuscript (version A) (see Figure 1). Version B offers additional flexibility to use BMD in a researcher’s desired output space and ROI format. In brief, version B is preprocessed in five output spaces (MNI152NLin2009cAsym, anatomical, fsaverage, fsnative, fsLR32k), contains beta estimates computed with GLMsingle (Prince et al., 2022) for MNI152NLin2009cAsym and fsLR32k spaces, and defines 47 ROIs in MNI152NLin2009cAsym space (left and right hemispheres of the 22 ROIs defined in the main text and MT, plus one “BMDgeneral” ROI). We show whole brain noiseceiling reliability results in the volume-based MNI152NLin2009cAsym and surface-based fsLR32k spaces and high predictivity of a motion energy model in motion-selective ROIs (MT, hV4, V3AB, IPS0).

Both version A (presented in the manuscript) and version B (described here) are identical up to fMRIPrep preprocessing (Esteban et al., 2019). Details on the experimental design, participants, and MRI acquisition protocols can be found in the main text. Version A was preprocessed using the default 6 degrees of freedom for BOLD to T1w image registration (the flag –bold2t1w-dof) and one standard volumetric output space (MNI152NLin2009cAsym). Version B was preprocessed with 12 degrees of freedom for BOLD to T1w image registration and four output spaces comprising a standard and native volume output (MNI152NLin2009cAsym, anat) and a standard and native surface output (fsaverage, fsnative) with transformation matrices available between the spaces (see fMRIPrep preprocessing boilerplate text below). We then use version B’s preprocessed results to convert the data to a fifth output space (fsLR32k) using Ciftify (Dickie et al., 2019). Made for the Human Connectome Project (Glasser et al., 2013; Van Essen et al., 2013), CIFTI format organizes brain data in 2D surface-based “grayordinates” for cortical structures and 3D volume-based voxels for subcortical structures. Ciftify adapts the HCP preprocessing pipeline to provide excellent volume-to-surface registration, especially for inter-subject analyses, and access to a suite of HCP analysis and visualization tools (Dickie et al., 2019; Glasser et al., 2013; Robinson et al., 2018).

We provide single trial beta estimates using GLMsingle (Prince et al., 2022) in the volume-based MNI152NLin2009cAsym and surface-based fsLR32k output spaces. We also define 47 ROIs in the MNI152NLin2009cAsym volume space for greater research and modeling flexibility. The 47 ROIs are similar to the 22 ROIs described in the main manuscript but are separated by left/right hemisphere, include the motion-selective MT ROI, include a “BMDgeneral” ROI that broadly defines reliably activated cortex across all BMD subjects, and enforce ROIs to have an equal number of voxels across subjects to facilitate inter-subject modeling.

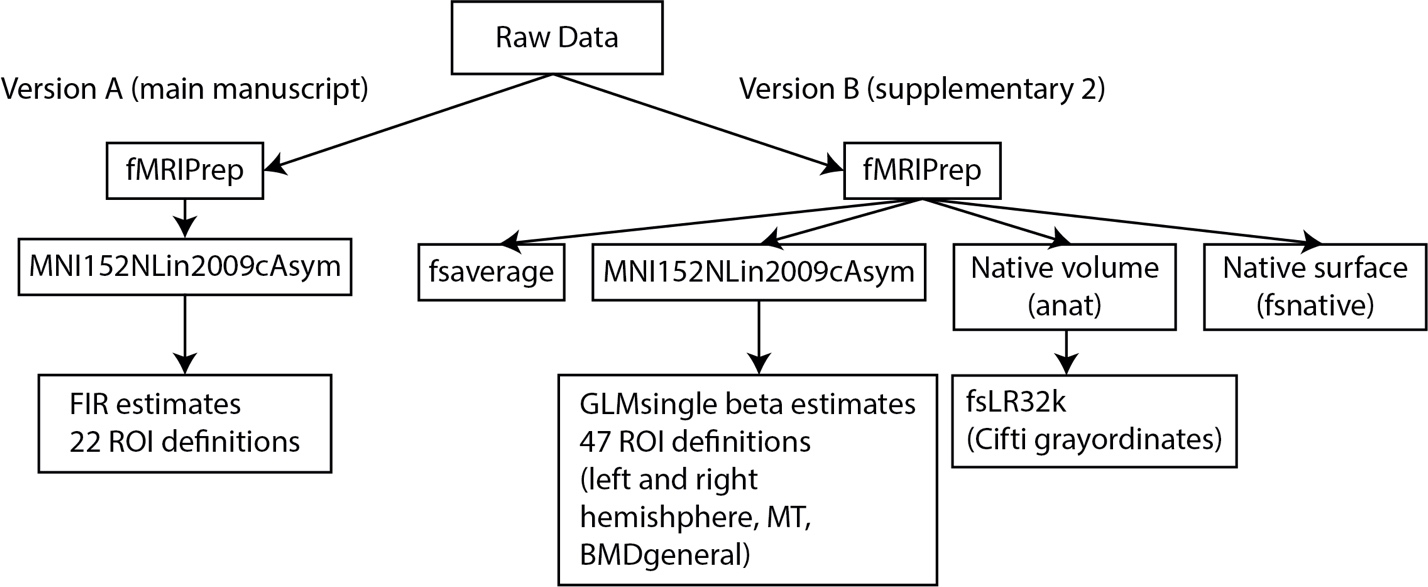

**Figure 1: Overview of preprocessing pipelines.** Version A (left) of BMD was preprocessed with fMRIPrep into a standard volumetric output space, modeled with FIR functions, and supplemented with 22 ROI definitions. Details are provided in the main manuscript. Version B (right) of BMD was preprocessed with fMRIPrep into two volume-based and two surface-based output spaces. Single trial beta estimates using GLMsingle and 47 ROI definitions were provided in the standard volume-based space. Data was transformed into a fifth output space (fsLR32k) using the Ciftify toolbox and scripts from the Human Connectome Project preprocessing pipeline.

2 Version B MNI152NLin2009cAsym Preprocessing and ROI Definition

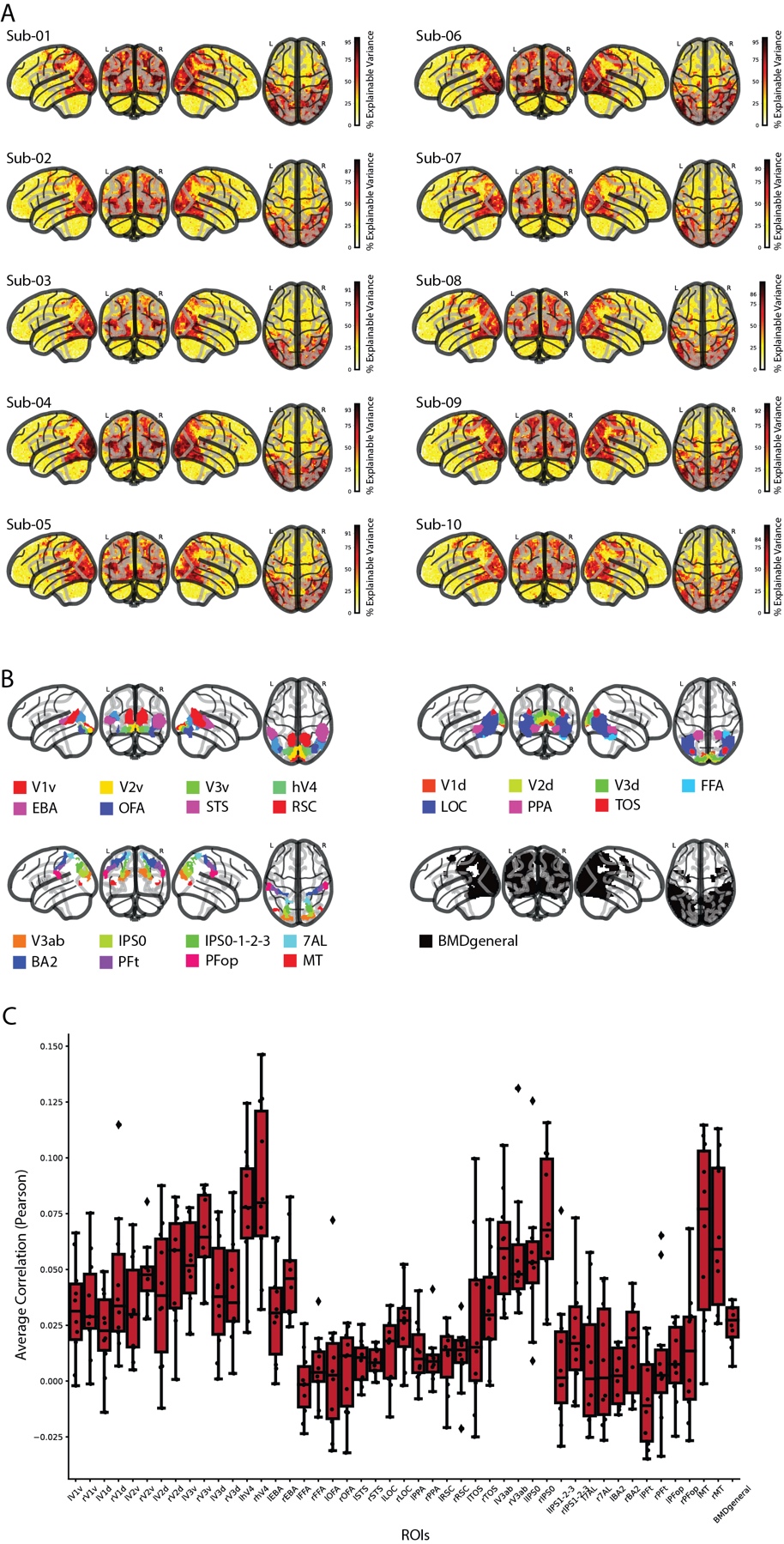

**Figure 2: Whole-brain noiseceiling and regions of interest (ROIs)** A) For each subject and voxel in the whole brain, we show the noiseceiling as percent of explainable variance using the testing set videos. The colorbar in each plot is normalized between 0 and 100, and the highest colorbar tick is that subject's maximum explainable variance in the whole brain. B) We show the 46 non-overlapping parcels (combined left and right hemispheres) and BMDgeneral (black) in a glass brain. Each subject's 8 category-selective ROIs (EBA, OFA, STS, RSC, FFA, LOC, PPA, and TOS) are functionally defined by extracting the top 50% most active voxels within the respective parcel. All subjects share the same ROI definition from the remaining parcels. BMDgeneral is defined independently from the ROIs and reflects a group-averaged region of cortex that reliably responds to videos in the BOLD Moments experiment. C) The boxplots depict the correlation (Pearson) between the predicted brain responses with the true responses of the testing set using stimulus features computed from a motion energy model. The boxes show the median response across subjects (horizontal line), 25^th^ and 75^th^ percentile (lower and upper box boundary), and whiskers extending to maximum and minimum values within 1.5 times the interquartile range. Individual subject results are shown as black points, and outliers are shown as diamonds (n=10 subjects for all ROIs).

All fMRI data were organized in the standardized BIDS format (K. J. Gorgolewski et al., 2016) and preprocessed using fMRIPrep (Esteban et al., 2019). The data were slice time corrected, co-registered to the subject's T1w anatomical scan, and spatially normalized to a standard volumetric MNI152NLin2009cAsym template brain. The main experimental runs were then temporally interpolated from their acquisition TR of 1.75 seconds to a TR of 1 second to time-lock volume sampling to stimulus presentations. A General Linear Model (GLM) was used to estimate single trial beta estimates (Prince et al., 2022). The beta responses were then z-scored across video conditions.

2.1 General Linear Model

2.1.1 Functional Localizer

We use GLMsingle (Prince et al., 2022) to model the hemodynamic response function (HRF) to the video localizer for each subject separately. The subject’s preprocessed data in MNI152NLin2009cAsym space was spatially smoothed with a 9mm full width half maximum of the gaussian kernel. The data was then temporally interpolated from an acquisition TR of 1.75s to an interpolated TR of 1s to timelock image acquisition to block onset. Each block, although composed of 6 3s videos (except for the fixation blocks, where no videos were shown), was modeled as a single stimulus. The onsets and durations (18s) of the Body, Face, Object, Scene, Scrambled, and Fixation blocks, along with the temporally interpolated and smoothed fMRI time series, were input to the general linear model. GLMsingle (1) chose an optimal HRF from a library at each voxel, (2) identified a number of nuisance regressors from principal component analysis of a noise pool that explain a maximum amount of variance, and (3) performed fractional ridge regression at each voxel to estimate single trial betas.

2.1.2 Main Experiment

For each subject, we fit beta estimates to each single-trial fMRI response in the main experiment using GLMsingle (Prince et al., 2022). The preprocessed data in MNI152NLin2009cAsym space was temporally interpolated from an acquisition TR of 1.75s to an interpolated TR of 1s to time-lock stimulus onset to image acquisition (e.g., 1.75s does not evenly go into the inter-trial interval of 4s). In this way, we acquire fMRI scans at different timepoints along the BOLD signal (with respect to stimulus presentation) and, after interpolating, achieve a regular sampling of the BOLD signal time-locked to stimulus onset for easier analysis. The interpolated fMRI time series, stimuli onsets, and stimuli durations (modeled as a 0s impulse) for each session separately were input to the general linear model. GLMsingle estimated single trial beta values by (1) fitting an optimal HRF to each voxel from a library of HRFs, (2) identifying nuisance regressors from a noise pool that maximally explain variance, and (3) implementing fractional ridge regression to improve estimates in a rapid event-related design. Responses to testing and training videos were estimated separately.

In this way, we obtained one beta estimate for each stimulus presentation for each subject. This resulted in a total of 4,020 beta estimates per subject (3 beta estimates x 1,000 training videos and 10 beta estimates x 102 testing videos).

Nan-indices corresponding to outside the subject’s brain mask were identified and removed. For the training and testing data separately, the beta estimates were then z-scored across video conditions such that the response profile for each stimulus presentation at each voxel (i.e., a vector of length 1000 for training data or of length 102 for testing data) had a mean value of 0 and standard deviation of 1.

2.2 Regions of Interest Definition

We computed a non-overlapping set of 46 ROIs (regions of interest) (23 ROIs separated by left and right hemispheres) previously known to be driven by dynamic stimuli spanning visual and parietal cortices (Gazzola & Keysers, 2009; Le et al., 2017; Logothetis & Sheinberg, 1996; R. Peeters et al., 2009; R. R. Peeters et al., 2013; Rizzolatti & Sinigaglia, 2010; Silver & Kastner, 2009; VanRullen & Thorpe, 2001). Note that these ROI definitions differ slightly compared to those detailed in the main manuscript (version A). We first created a non-overlapping parcellation in the standard MNI152NLin2009cAsym space identical across subjects composed of parcels resampled from Wang and colleagues, Glasser and colleagues, and Julian and colleagues (Glasser et al., 2016; Julian et al., 2012; Wang et al., 2015). Finally, we used the t-contrasts from each subject’s functional localizer results to identify the top 50% of voxels within the corresponding functional parcel from Julian and colleagues (Julian et al., 2012) (bodies > objects: EBA; objects > scrambled: LOC; scenes > objects: PPA, RSC, STS; faces > objects: OFA, FFA, STS). This ROI definition method facilitated inter-subject modeling approaches by ensuring all ROIs were defined for each subject and each ROI contained the same number of voxels across subjects. Furthermore, the parcellation shared across subjects (before taking each subject’s top 50% of voxels in the parcels from Julian and colleagues (Julian et al., 2012)) allowed modeling approaches that incorporate voxel-level spatial information, since the parcel indices are the same for each subject.

In detail, we defined ROIs V1v, V1d, V2v, V2d, V3v, V3d, hV4, V3a, V3b, IPS0, IPS1, IPS2, IPS3 from Wang and colleagues (Wang et al., 2015) (maxprob_vol_{h}h.nii, where {h} is “l” or “r”), L_2 (here referred to as BA2), 7AL, PFt, PFop, and MT from Glasser and colleagues (Glasser et al., 2016) (HCPMMP1_on_MNI152_ICBM2009a_nlin.nii), and EBA, LOC, PPA, RSC, STS, OFA, FFA, and STS from Julian and colleagues (Julian et al., 2012) ({h}{parcel}.img from the n=30 group, where {h} is “l” or “r” and {parcel} is the parcel name). We group V3a and V3b into V3ab and IPS1, IPS2, and IPS3 into IPS1-2-3 due to subtle differences in functional preferences that can be difficult to resolve with our in-the-wild naturalistic stimuli (Georgieva et al., 2009; Konen & Kastner, 2008; Press et al., 2001; Smith et al., 1998). All ROIs were resampled into our functional volumetric dimensions and separated by left and right hemispheres. Voxels outside a common brain mask computed across subjects were removed from the parcel. There was no overlap between the parcels within Wang and colleagues (Wang et al., 2015) or between the parcels within Glasser and colleagues (Glasser et al., 2016).

To address minimal overlap between the parcels derived from Julian and colleagues (Julian et al., 2012), the in-question voxels were assigned to the parcels that had a greater inter-subject agreement from our functional localizer experiment. Specifically, a t-test (two-sided, independent) was computed between the condition beta estimates to define category-selective contrasts: bodies > objects (body selective; EBA), objects > scrambled (object selective; LOC), scenes > objects (scene selective; PPA, RSC, TOS), and faces > objects (face selective; OFA, FFA, STS). Similar to the Group-constrained Subject Specific (GSS) procedure (Julian et al., 2012), the t-contrast maps were binarized at a p-value cutoff (p < 0.05, uncorrected), where voxels below this cutoff were assigned 0 and voxels above this cutoff were assigned 1. The binarized t-contrast maps were averaged across subjects and smoothed (6mm full width half maximum of the gaussian kernel) to obtain four probability maps (one for each contrast) that contains information on inter-subject agreement at each voxel. If both LOC and EBA overlapped at voxel A, for example, we indexed voxel A’s value in both the objects > scrambled probability map (for LOC) and the bodies > objects probability map (for EBA) and assigned voxel A to the parcel with the higher value (i.e., the higher inter-subject agreement). No parcels within a contrast overlapped.

Finally, we addressed the minimal overlap between parcels across the three atlases. If non-functionally defined parcels (i.e., the parcels derived from Wang and colleagues (Wang et al., 2015) and Glasser and colleagues (Glasser et al., 2016)) overlapped with functionally-defined parcels (i.e., from Julian and colleagues (Julian et al., 2012)), preference was given first to the non-functionally defined parcel. Otherwise, the voxel may end up not being assigned to any ROI after subject-specific ROI definition (described below) because the functionally-defined parcels are generous in size, reflecting group-level inter-subject agreement. If the overlapping parcels were all non-functionally defined, preference would have gone to the smaller parcel to preserve its size, but there was only overlap between functionally and non-functionally defined parcels. In this way, we obtained 46 non-overlapping parcels identical for each subject.

We then functionally define the category-selective ROIs for each subject by identifying the top 50% most active (i.e., highest t-values, uncorrected) voxels inside the ROIs respective t-contrast map masked by the corresponding parcel (Mineroff et al., 2018). This method achieves both a subject-specific functional definition, maintains the relative size between ROIs, and ensures the same number of voxels within an ROI across subjects. No constraint of contiguity was enforced.

We additionally define a swathe of cortex that showed consistently reliable responses to videos across subjects in this study, here termed BMDgeneral. BMDgeneral was algorithmically defined in five steps: (1) compute split-half correlations across stimuli repetitions to obtain p-values at each voxel, (2) binarize the volume at p < 0.05 (uncorrected), indicating voxels with a value of 0 and 1 have poor and good split-half reliability, (3) average each subject's binarized mask to obtain a probability map reflecting the inter-subject agreement of each voxel's reliability, (4) smooth the probability map (6mm at full width half maximum), and (5) identify clusters (cluster threshold = 50 voxels, 8mm between peaks, statistic threshold of 0.1). All identified clusters (here, 4 clusters) are collectively identified as BMDgeneral. BMDgeneral may or may not overlap with the 46 ROIs.

3 Version B CIFTI Preprocessing

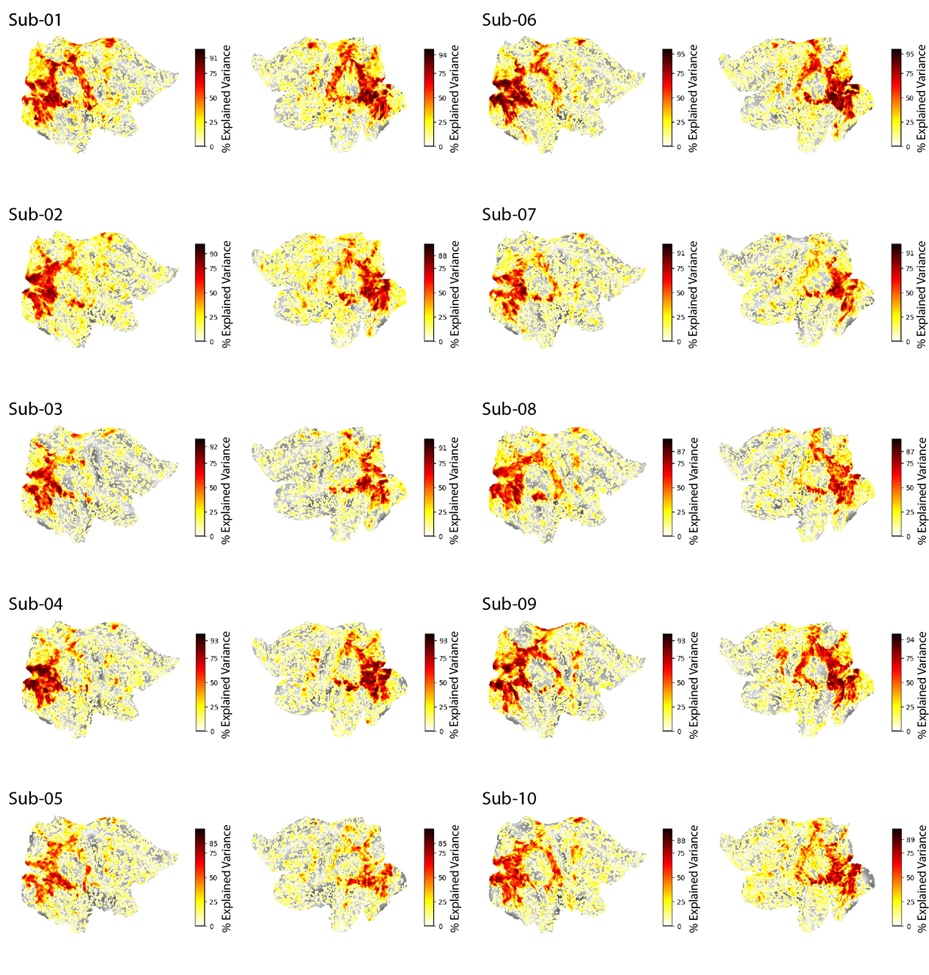

**Figure 3: Left and right cortex noiseceiling.** For each subject and grayordinate vertex in the left and right flattened hemispheres, we show the noiseceiling as percent of explainable variance using the testing set videos. The colorbar in each plot is normalized between 0 and 100, and the highest colorbar tick is that subject's maximum explainable variance in the whole brain. Values are thresholded at 1.

After fMRIPrep preprocessing, data was converted to CIFTI format using the Ciftify framework (version 2.3.3) (Dickie et al., 2019), which adapts scripts from the Human Connectome Project (HCP) preprocessing pipeline (Glasser et al., 2013). Ciftify allows registration of data to fsLR32k output space in a HCP compatible directory structure while still respecting BIDS organization (K. J. Gorgolewski et al., 2016)for the use of BIDS-apps (e.g., fMRIPrep). First, the “ciftify_recon_all” function converts a subject’s freesurfer output directory (output from fMRIPrep) into CIFTI format, registers their anatomical data from to a surface-based mesh using the MSMSulc algorithm (Robinson et al., 2018), then resamples to a 32k standard space (fsLR32k). Second, the “cifity_subject_fmri” maps the preprocessed functional scans to the subject’s fsLR32k space. Voxels along the cortical ribbon are weighted and projected to vertices on the surface (e.g., “grayordinates”), and subcortical voxels (including cerebellum) are resampled to volumetric MNI space. No smoothing was applied.

Note that preprocessing BMD through the entire HCP preprocessing pipeline is expected to obtain even better results due to the availability of high resolution T2w and fieldmap scans. We use Ciftify here to maintain a BIDS compliant dataset (for use with other BIDS-apps) and a common fMRIPrep preprocessing root between output spaces all while still making the data immediately available in the advantageous CIFTI format.

3.1 General Linear Model

3.1.1 Main Experiment

We estimate single-trial beta values at each vertex on the fsLR32k cortical surface for each subject using GLMsingle (Prince et al., 2022). The unsmoothed time series output from Ciftify (Dickie et al., 2019) was temporally interpolated to a TR of 1s (acquisition TR=1.75s) in order to temporally align image acquisition with stimulus onset. The video stimulus trials were entered into the general linear model (GLM) as regressors of interest. Null trials were not entered into the GLM. GLMsingle then computed single-trial beta estimates by optimally fitting one of twenty hemodynamic response functions (HRFs) to each voxel, calculating nuisance regressors from noisy vertices, and incorporating fractional ridge regression for improved beta estimates in a rapid event-related experimental designs. The GLM was computed per session, and testing and training runs were computed independently. This procedure resulted in 4,020 single-trial beta estimates for each subject.

In this way, we obtained one beta estimate for each stimulus presentation for each subject. This resulted in a total of 4,020 beta estimates per subject (3 beta estimates x 1,000 training videos and 10 beta estimates x 102 testing videos).

4 Noiseceiling Calculation

We compute the percent of explainable variance at each voxel for each subject as an estimate of the noiseceiling (equation 2) (Allen et al., 2022; Prince et al., 2022). First, the noise, signal and total variance of the beta estimates is computed. The noise variance is computed as the mean variance of the beta estimates of the within-video presentation trials. The total variance is computed as the variance of the beta estimates across all video presentation trials. The signal variance is computed as the total variance minus the noise variance. The signal variance was positively rectified, where negative values were assigned 0 and positive values were preserved. Next, the noiseceiling signal-to-noise ratio (SNR) (*ncsnr*) is computed as the fraction of signal standard deviation ($\sigma_{signal}$) to noise standard deviation ($\sigma_{noise}$) (equation 1),

$ncsnr=\frac{\sqrt{{var}_{signal}}}{\sqrt{{var}_{noise}}}$ (equation 1)

where the standard deviation is equal to the square root of the variance ($\sigma=\sqrt{var}$). Finally, the percentage of explainable variance (*noiseceiling*) is calculated as,

$noiseceiling=\frac{{ncsnr}^{2}}{{ncsnr}^{2}+\frac{1}{numTrials}}*100$ (equation 2)

where *ncsnr* is the noiseceiling signal-to-noise ratio (SNR) and *numTrials* is the number of video presentation trials (*numTrials*=10 for the testing set and *numTrials*=3 for the training set). This measure of voxel reliability differs from the split-half reliability measure used in the main manuscript (version A) but produce very similar results. The reliability measure proposed here is computationally less expensive than a split-half computation and has no stochastic elements.

5 Motion Energy Features Computation and Encoding Model

Motion energy features were used to predict brain activity in response to BMD’s 3 second naturalistic videos. The motion energy model (Adelson & Bergen, 1985; Nishimoto et al., 2011; Watson & Ahumada, 1985) consists of a series of spatial and temporal Gabor filters intended to capture local motion and direction in a video stimulus, thus making it a highly interpretable method to model video dynamics. The motion energy encoding model accuracy (Figure 2C) shows highest prediction accuracy in motion selective ROIs, namely MT (Born & Bradley, 2005; Nishimoto & Gallant, 2011), hV4 (Kamitani & Tong, 2006; Roe et al., 2012), V3AB (Konen & Kastner, 2008; Smith et al., 1998), and IPS0 (Konen & Kastner, 2008). These results support that single trial beta estimates of BMD’s 3 second naturalistic videos capture motion information.

Motion energy features for each BMD video stimulus was computed using the MATLAB code available here: <https://github.com/gallantlab/motion_energy_matlab> (Nishimoto et al., 2011; Nishimoto & Gallant, 2011). For each 268x268 video, the frames were converted from RGB to LAB color space, and only the L (luminance) channel was retained. The luminance channel was then passed through a three-dimensional bank of spatiotemporal Gabor filters consisting of two spatial dimensions and one temporal dimension. Similar to the filter bank used in (Nishimoto et al., 2011) to model naturalistic movies, the three-dimensional filters are defined at five spatial frequencies (0, 2, 4, 8, 16, and 32 cycles/image), three temporal frequencies (0, 2, and 4Hz), and eight directions (0, 45, 90, 135, 180, 225, 270, and 315 degrees) with the exception that the 0 Hz temporal filter is defined at only 0, 45, 90, and 135 degrees directions and the 0 cycles/image spatial filter is defined at 0 degree orientation. Local motion-energy features were computed by taking the square root of the sum of the squared outputs of each pair of filters with orthogonal phases. The logarithm of the output from these filters was computed to scale large values, and the temporal dimension of the output was downsampled to 1 second to match the fMRI sampling rate (i.e., the interpolated TR of 1 second) of the BOLD time series. The output was then z-scored across time. In total, this procedure resulted in a matrix of size 3 x 6555 (seconds x motion energy features).

The motion energy features were then used in a voxelwise linear encoding model (Naselaris et al., 2011) to predict the brain activity (beta estimates) in 47 regions of interest (ROIs) from the version B preprocessed data in MNI152NLin2009cAsym space (Figure 2C). Specifically, the motion energy features for each video were concatenated along the three seconds and underwent principal component analysis (PCA) to reduce dimensionality to the top 100 components. PCA was fit to the training videos and applied to both the training and testing videos. A linear model was then fit to the training video features to predict the response at the voxel. The learned weights of the linear model were then applied to the testing video features. The encoding model accuracy was computed as the correlation of the vector of predicted responses of the test set with the vector of true responses of the test set.

6 fMRIPrep Preprocessing Boilerplate Text

We reproduce the fMRIPrep boilerplate text describing version B’s preprocessing details below:

“”

Results included in this manuscript come from preprocessing performed using fMRIPrep 23.0.2 (Esteban et al., 2019, 2022; RRID:SCR_016216), which is based on Nipype 1.8.6 (Esteban, Oscar et al., 2022; K. Gorgolewski et al., 2011; RRID:SCR_002502).

#### Preprocessing of B0 inhomogeneity mappings

A total of 6 fieldmaps were found available within the input BIDS structure for this particular subject. A B0 nonuniformity map (or fieldmap) was estimated from the phase-drift map(s) measure with two consecutive GRE (gradient-recalled echo) acquisitions. The corresponding phase-map(s) were phase-unwrapped with prelude (FSL 6.0.5.1:57b01774).

#### Anatomical data preprocessing

A total of 1 T1-weighted (T1w) images were found within the input BIDS dataset. The T1-weighted (T1w) image was corrected for intensity non-uniformity (INU) with N4BiasFieldCorrection (Tustison et al., 2010), distributed with ANTs 2.3.3 (Avants et al., 2008; RRID:SCR_004757), and used as T1w-reference throughout the workflow. The T1w-reference was then skull-stripped with a Nipype implementation of the antsBrainExtraction.sh workflow (from ANTs), using OASIS30ANTs as target template. Brain tissue segmentation of cerebrospinal fluid (CSF), white-matter (WM) and gray-matter (GM) was performed on the brain-extracted T1w using fast (6.0.5.1:57b01774, RRID:SCR_002823, Zhang et al., 2001). Brain surfaces were reconstructed using recon-all (FreeSurfer 7.3.2, RRID:SCR_001847, Dale et al., 1999), and the brain mask estimated previously was refined with a custom variation of the method to reconcile ANTs-derived and FreeSurfer-derived segmentations of the cortical gray-matter of Mindboggle (RRID:SCR_002438, Klein et al., 2017). Volume-based spatial normalization to one standard space (MNI152NLin2009cAsym) was performed through nonlinear registration with antsRegistration (ANTs 2.3.3), using brain-extracted versions of both T1w reference and the T1w template. The following template was were selected for spatial normalization and accessed with TemplateFlow (23.0.0, Ciric et al., 2022): ICBM 152 Nonlinear Asymmetrical template version 2009c [Fonov et al., 2009, RRID:SCR_008796; TemplateFlow ID: MNI152NLin2009cAsym].

#### Functional data preprocessing

For each of the 62 BOLD runs found per subject (across all tasks and sessions), the following preprocessing was performed. First, a reference volume and its skull-stripped version were generated using a custom methodology of *fMRIPrep*. Head-motion parameters with respect to the BOLD reference (transformation matrices, and six corresponding rotation and translation parameters) are estimated before any spatiotemporal filtering using mcflirt (FSL 6.0.5.1:57b01774, Jenkinson et al., 2002). The estimated fieldmap was then aligned with rigid-registration to the target EPI (echo-planar imaging) reference run. The field coefficients were mapped on to the reference EPI using the transform. BOLD runs were slice-time corrected to 0.834s (0.5 of slice acquisition range 0s-1.67s) using 3dTshift from AFNI (Cox & Hyde, 1997, RRID:SCR_005927). The BOLD reference was then co-registered to the T1w reference using bbregister (FreeSurfer) which implements boundary-based registration (Greve & Fischl, 2009). Co-registration was configured with twelve degrees of freedom to account for distortions remaining in the BOLD reference. Several confounding time-series were calculated based on the *preprocessed BOLD*: framewise displacement (FD), DVARS and three region-wise global signals. FD was computed using two formulations following Power (absolute sum of relative motions, Power et al., 2014) and Jenkinson (relative root mean square displacement between affines, Jenkinson et al., 2002). FD and DVARS are calculated for each functional run, both using their implementations in *Nipype* (following the definitions by Power et al., 2014). The three global signals are extracted within the CSF, the WM, and the whole-brain masks. Additionally, a set of physiological regressors were extracted to allow for component-based noise correction (*CompCor*, Behzadi et al., 2007). Principal components are estimated after high-pass filtering the *preprocessed BOLD* time-series (using a discrete cosine filter with 128s cut-off) for the two *CompCor* variants: temporal (tCompCor) and anatomical (aCompCor). tCompCor components are then calculated from the top 2% variable voxels within the brain mask. For aCompCor, three probabilistic masks (CSF, WM and combined CSF+WM) are generated in anatomical space. The implementation differs from that of Behzadi et al. in that instead of eroding the masks by 2 pixels on BOLD space, a mask of pixels that likely contain a volume fraction of GM is subtracted from the aCompCor masks. This mask is obtained by dilating a GM mask extracted from the FreeSurfer’s aseg segmentation, and it ensures components are not extracted from voxels containing a minimal fraction of GM. Finally, these masks are resampled into BOLD space and binarized by thresholding at 0.99 (as in the original implementation). Components are also calculated separately within the WM and CSF masks. For each CompCor decomposition, the k components with the largest singular values are retained, such that the retained components’ time series are sufficient to explain 50 percent of variance across the nuisance mask (CSF, WM, combined, or temporal). The remaining components are dropped from consideration. The head-motion estimates calculated in the correction step were also placed within the corresponding confounds file. The confound time series derived from head motion estimates and global signals were expanded with the inclusion of temporal derivatives and quadratic terms for each (Satterthwaite et al., 2013). Frames that exceeded a threshold of 0.5 mm FD or 1.5 standardized DVARS were annotated as motion outliers. Additional nuisance timeseries are calculated by means of principal components analysis of the signal found within a thin band (crown) of voxels around the edge of the brain, as proposed by (Patriat et al., 2017). All resamplings can be performed with a single interpolation step by composing all the pertinent transformations (i.e. head-motion transform matrices, susceptibility distortion correction when available, and co-registrations to anatomical and output spaces). Gridded (volumetric) resamplings were performed using antsApplyTransforms (ANTs), configured with Lanczos interpolation to minimize the smoothing effects of other kernels (Lanczos, 1964). Non-gridded (surface) resamplings were performed using mri_vol2surf (FreeSurfer).

Many internal operations of *fMRIPrep* use *Nilearn* 0.9.1 (Abraham et al., 2014, RRID:SCR_001362), mostly within the functional processing workflow. For more details of the pipeline, see [the section corresponding to workflows in *fMRIPrep*’s documentation](https://fmriprep.readthedocs.io/en/latest/workflows.html).

“”
